# Mechanistic interpretation of non-coding variants helps discover transcriptional regulators of drug response

**DOI:** 10.1101/503458

**Authors:** Xiaoman Xie, Casey Hanson, Saurabh Sinha

## Abstract

Identification of functional non-coding variants (polymorphisms) and their mechanistic interpretation is a major challenge of modern genomics, especially for precision medicine. Transcription factor (TF) binding profiles and epigenomic landscapes in reference samples can help us functionally annotate the genome, but do not provide ready answers regarding the effects of non-coding variants. A promising computational approach is to build models that predict TF-DNA binding from sequence, and use such models to score a variant’s impact on TF binding strength. Here, we asked if this mechanistic approach to variant interpretation can be combined with information on genotype-phenotype associations to discover important transcription factors regulating phenotypic variation among individuals. We developed a statistical approach that integrates phenotype, genotype, gene expression, TF ChIP-seq and Hi-C chromatin interaction data to answer this question. Using drug sensitivity measured in lymphoblastoid cell lines as the phenotype of interest, we tested if the non-coding variants statistically linked to the phenotype are enriched for strong predicted impact on DNA-binding strength of a TF, and used this test to identify TFs regulating individual differences in the phenotype. Our method relies on a new method for predicting variant impact on TF-DNA binding, that uses a combination of biophysical modelling and machine learning. We report statistical and literature-based support for many of the TFs discovered here as regulators of drug response variation. We show that the use of mechanistically driven variant impact predictors can identify TF-drug associations that would otherwise be missed. We examined in depth the evidence underlying one reported association – that of the transcription factor ELF1 with the drug doxorubicin – and identified several genes that may mediate this regulatory relationship.

## INTRODUCTION

A major open problem today is how differences in DNA sequence, e.g., single-nucleotide polymorphisms (SNPs) and variants (SNVs), lead to health-related and other phenotypic differences among individuals. A common approach is to find polymorphisms/variants that are statistically correlated with phenotypic differences, as in genome-wide association studies (GWAS) (1), family-based association tests (2), and expression quantitative trait loci (eQTLs) (3,4) for phenotype-related genes. However, statistically identified variants may not be functionally related to phenotypes (5), due to a variety of factors including linkage disequilibrium (LD). This problem is particularly pronounced in the case of non-coding variants, which represent the vast majority of GWAS findings (6,7) and often function by influencing gene regulation. Accurate contextual information about non-coding variants can improve our ability to disambiguate variants causally related to gene expression and phenotype (8,9) from nearby non-functional SNPs. For example, if we have prior knowledge of a relevant transcription factor (TF), then the presence of a variant within a TF binding site (TFBS) may add to our confidence in the variant’s regulatory potential; the assumption here is that such a variant influences the TF’s binding to that site and consequently the gene regulatory impact of the TF. Advanced techniques for predicting in vivo TF-DNA binding potential from DNA sequence (gkm-SVM (10,11), DEEP-BIND (12), DEEPSEA (13), DeFine (14), and Sasquatch (15)) can facilitate this approach by providing more accurate estimates of a variant’s impact on TF binding. In addition to providing a means for statistically prioritizing those non-coding variants by their likelihood of functionality, this strategy also offers a mechanistic explanation about their function, i.e., their impact on the TF-gene regulatory relationship. For example, Zhang et al. (5) adopted such a strategy: they combined a method for predicting changes in TF binding with multi-omics data to identify a SNP that impacts the binding strength of a TF called GATA3 to modulate breast cancer susceptibility.

It must be noted, however, that the above approach to identify phenotype-related non-coding variants along with their regulatory mechanism is still in its infancy and its sensitivity-specificity tradeoff is not well understood. Reliable mechanistic claims of a SNP mediating a TF’s influence on phenotypic variation often require multiple lines of ‘-omic’ evidence as well as prior knowledge. A related but less explored opportunity is to examine a collection of variants associated with a phenotype (e.g., from a GWAS study) and test the collection for enrichment of variants predicted to impact TF-DNA binding; such an enrichment can associate the TF, rather than individual variants, with the phenotype. This may give us mechanistic insights of a more global nature, with greater confidence than what the available data allows at the level of individual SNPs. In recent work, we adopted this general strategy to identify transcription factors regulating phenotypic variation across individuals, by combining genotype, gene expression and phenotype information with genome-wide profiles of TF-DNA binding. The underlying principles were twofold: (1) If a gene’s expression is correlated with phenotype, and a SNP correlated with that gene’s expression (eQTL of the gene) is located within a TFBS, we treated this as (weak) evidence that the TF influences the phenotype via that gene; (2) if such evidence is observed significantly many times, i.e., across many genes, we hypothesized that the TF plays an important regulatory role in phenotypic variation. The assumption is that TF binding is affected by the SNP and this effect underlies the SNP’s correlation with gene expression, which in turn contributes to phenotypic variation. We pursued this line of reasoning in (16,17) to systematically identify, through statistical testing and probabilistic graphical models, major TFs associated with a specific type of phenotypic variation, viz., differences in cytotoxic response to a particular drug in a panel of cell lines. Our goal in the current work is to test if information about the functional impact of variants on TF-binding can improve inferences of TF-phenotype associations.

Two of the three pieces of information considered in the above scheme – (a) strength of SNP association with gene expression (eQTL) and (b) gene expression correlation with phenotype (a transcriptome wide association study or ‘TWAS’ (18)) – are quantified by relatively established procedures. However, the third axis of information crucial to the approach – the evidence that a TF’s binding, and hence its regulatory influence on a gene, is affected by a SNP – is harder to assess. In previous studies, we treated the presence of a SNP inside a ChIP peak of the TF, located within the 50 Kbp upstream region of the gene, as such evidence. However, this heuristic has obvious limitations. First, a SNP located within a ChIP peak may not necessarily impact the TF’s binding. This may be addressed by borrowing ideas from previous studies (19,20) that have used motif and k-mer based scans within ChIP peaks to identify regulatory SNPs likely to affect that TF’s binding. Second, a TF binding event located further than 50 Kbp from the TSS may also exert regulatory influence on a gene, depending on chromatin looping structures (21); conversely, every TF binding event located within a modest distance (e.g., 50 Kbp) of the TSS does not necessarily have a regulatory influence on the gene. Use of chromatin interaction data sets offers a resolution of this issue (21). In this work, we address the above limitations of ascribing a regulatory relationship to a (TF, SNP, gene) triplet, through a combination of established and novel methods, with the express goal of aggregating such evidences and combining them with gene-phenotype correlations to discover regulatory mechanisms underlying phenotype variation.

We develop and use a new computational pipeline to identify TFs associated with drug response variation across individuals, building on the ideas outlined above, and performing integrative analysis of genotype, gene expression and cytotoxicity data on a panel of ~300 cell lines, along with TF-ChIP data from ENCODE and TF binding motifs from various databases. We utilize a state-of-the-art, ‘k-mer’ based, machine learning technique to predict the impact of a SNP on TF binding strength. We also develop an alternative method for this task, which uses one or more motifs known to represent a TF’s binding preferences, and combines biophysically inspired modeling and machine learning ideas. Through systematic benchmarking, we find that this motif-based method is at least as effective as the ‘k-mer’ based technique for predicting allele-specific TF-DNA binding, in contrast to recent reports that leading k-mer based approaches clearly outperform motif-based approaches (22). Ultimately, using both k-mer based and motif-based predictors, and utilizing chromatin interaction domains and loops to link variants to genes, we show that modern tools of SNP impact prediction can lead to the discovery of novel regulatory mechanisms underlying phenotypic variation that are missed when not using SNP impact predictors. By aggregating evidence from many SNPs with putative effects on TF binding, we systematically identify TFs that influence individual-level differences in drug sensitivity, for several cytotoxic drugs. We examine one such discovered association more closely, viz., the predicted and experimentally confirmed effect of the TF ‘E74-like factor 1’ (ELF1) on sensitivity to the drug doxorubicin. Our analysis suggests several genes that may be under ELF1 regulation and related to the doxorubicin response pathway.

## MATERIAL AND METHODS

### Data collection

Genotype, gene expression and drug response data on 95 Han-Chinese, 96 Caucasian and 93 African-American lymphoblastoid cell lines (LCL) from the Coriell Cell Repository (Camden, NJ, USA), of which 176 were female and 104 male. 1,344,658 germline SNPs were genotyped and SNPs with minor allele frequency < 5% or which deviated from Hardy-Weinberg equilibrium < 95% were removed. Strand information was collected from dbSNP. 1,283,254 SNPs with same identifier and location in both LCL data and dbSNPs are used. Gene expression data consisted of 54,613 Affymetrix U133 Plus 2.0 Gene-ChIP (Santa Clara, CA, USA) probes assayed for the 284 individuals, with raw expression data being transformed using GC Robust Multi-Array Averaging (GC-RMA). Genotype and gene expression data are available at the National Center for Biotechnology Information (NCBI) Gene Expression Omnibus (http://www.ncbi.nlm.nih.gov/geo) under SuperSeries accession no. GSE24277. These data were published in a study by Niu et al. Gene mappings to the Affymetrix arrays were obtained for the Affymetrix Human Genome U133 Plus 2.0 array. ENSEMBL gene symbols were used as the gene reference of choice: we used 55,038 ENSEMBL gene symbols that were annotated with at least one ENSEMBL exon. Of the 54,613 probes assayed on the HG U133 Plus 2.0 array, 37,677 mapped to at least one of the 55,038 ENSEMBL gene symbols.

Drug response data were derived from dosage–response curves of 24 cytotoxic treatments published in Hanson et. al (17): 6MP, 6TG, ARAC, arsenic, carboplatin, CDDP, cladribine, docetaxel, doxorubicin, epirubicin, everolimus, fludarabine, gemcitabine, hypoxia, metformin, MPA, MTX, NAPQI, oxaliplatin, paclitaxel, radiation, rapamycin, TCN and TMZ. The phenotype, called EC50, represents the concentration at which the drug reduces the population of LCL cells to half of the initial population. Cytotoxicity assays were performed for every one of these drugs using the LCL panel. After initial optimization, cells were treated with a range of concentrations for any given drug tested, followed by incubation for 48 to 72 h. MTS cytotoxicity assays were then performed using Cell Titer 96 AQueous Non-Radioactive Cell Proliferation Assay kit (Promega Corporation, Madison, WI, USA), followed by absorbance measurement at 490 nm in a Safire2 microplate reader (Tecan AG, Switzerland). Cytotoxicity phenotypes were determined by the best fitting curve using the R package ‘drc’ (dose–response curve) (23) based on a logistic model.

### Transcription factor binding motifs

225 PWMs for 37 TFs were collected from three sources:

1. 29 PWMs for 27 TFs were collected from ENCODE factor book motifs from (http://hgdownload.soe.ucsc.edu/goldenPath/hg19/database/factorbookMotifPwm.txt.gz).
2. 185 PWMs based on ChIP data for 37 TFs from GM12878 cell line were downloaded from Factorbook website (http://www.factorbook.org/human/chipseq/tf/).
3. 25 PWMs for 21 TFs were obtained from HOCOMOCO Human (v10) (24), via the motif library of the MEME software (25).

All motifs are included as Supplementary File 1.

### ChIP-seq and accessibility data

We used ChIP-seq data from the ENCODE project, as summarized in the ‘Txn Factor’ track at the UCSC genome browser (‘wgEncodeRegTfbsClusteredWithCellsV3’ bed files). Clustered peaks observed in GM12878 cell line were used in this study. We also used genome-wide profiles of ChIP-seq signal values from ENCODE project (www.encodeproject.org). Signal values are used as numeric measurements of the TF binding strength for training and testing TF-DNA binding prediction. DNaseI hypersensitivity (DHS) uniform peaks for GM12878 cell line (ENCODE project) were downloaded from the UCSC web site.

### Training set generation

MOP, STAP and gkmSVM need to be trained on ChIP-seq data using DNA sequences and corresponding ChIP scores. For training purposes, we generated balanced training data sets for each TF, which composed of positive sequences and same number of negative sequences. We selected 1000 segments of 500bp length from each TF’s ChIP peaks as the positive set. (We limited the selection to peaks located within 50 Kbp upstream of a protein coding gene and excluded ‘High Occupancy Target’ or HOT regions, i.e., peaks overlapping 6 or more TFs with at least 50% overlap.) We defined a large collection of ‘negative windows’ for a TF to be 500 bp long segments in the positive sets of other TFs but not bound by the test TF. We then randomly selected 1000 windows from this collection as the negative set for the test TF. DNA sequence and signal value for each window was extracted from reference genome (hg19) and ChIP-seq data from ENCODE project. Thus a balanced dataset with 2000 windows was generated for each of the 37 TFs. These datasets were further separated into a balanced training set with 1600 windows and a balanced test set of 400 windows. (See Supplementary Note 1)

### Prediction of TF-DNA Binding

1. STAP: A separate STAP model (26) was trained for each of 225 PWMs (representing 37 distinct TFs) using the balanced training set. Cross validation (80% training, 20% testing) within these training data was used to learn a value for the site energy threshold (‘eT’) hyperparameter.
2. gkm-SVM: For each TF, a separate model was trained as recommended by authors (10,11), with default settings (http://www.beerlab.org/gkmsvm/gkmsvm-tutorial.htm).
3. MOP: The scores of a window reported by STAP models using different motifs of the TF were used as a feature vector representing the window and provided to a support vector machine (SVM). We trained an SVM model (package ‘e1071’ in R (27)) to predict ChIP scores from such feature vectors, using the same training data as above. Cross validation (80% training, 20% testing) within training data was used to learn hyperparameters.

To make the binding scores predicted by different methods fall on a comparable scale, we rescaled every score by a linear function so that the predicted binding scores for the 2000 windows in training and testing data range exactly from 0 to 1.

### Prediction of TFBS-SNP impact

We first generated a reference genome specific to our LCL genotype data set by starting with the ‘hg19’ reference genome and setting the nucleotide at each SNP location (in the LCL data set) to the major allele of that SNP in the data set. For each SNP, a 501bp window centered on that SNP was extracted from this LCL-specific reference genome, and two versions of its sequence – one with the minor allele and another with the major allele of that SNP – were used as inputs for TF binding predictors. The absolute value of the difference between predicted binding scores of these two sequences was used as the TFBS-SNP impact score (Delta-STAP, Delta-gkm-SVM or Delta-MOP, depending on the binding prediction method used). In this step, binding predictors trained on all 2000 windows defined above were used. The fourth method for SNP impact prediction, called Delta-PWM, directly uses the ‘Delta raw scores’ for ‘MEME signif PWM’ provided by Wagih et al (22).

### Evaluations on Allele-Specific-Binding (ASB) data

ASB SNPs and non-ASB SNPs for lymphoblastoid cell lines were collected from Wagih et al (22). Twenty-two of the 37 TFs, for which we have binding predictors, have ASB data for these cell lines. Among these 22 TFs, MEF2A, NFYB, SRF and USF1 have fewer than 150 annotated (ASB or non-ASB) SNPs, while SP1 and SPI1 did not have associated Delta-PWM data. For these reasons, these six TFs were excluded and we only used the data for the remaining 16 TFs in the ASB evaluation (Supplementary Table 2). The TFBS-SNP impact of each ASB and non-ASB SNP was measured by four methods (Delta-MOP, Delta-gkm-SVM, Delta-STAP and Delta-PWM) as explained above. AUROC and AUPRC values were calculated for each TF-method combination, indicating how well the corresponding impact score can be used to label the ASB and non-ASB SNPs.

### Identifying eQTLs in a gene’s regulatory region

We used Hi-C data (21) on 3-D chromatin architecture in the GM12878 cell line to construct the cis-regulatory region of each gene. First, the local ‘domain’ that the gene overlaps with was included in such a region. Second, for each pair of loci connected by a loop, if the gene overlaps with one of the loci, the other locus was included in its cis-regulatory region. For each SNP located within the cis-regulatory region of a gene, the association between genotype and gene expression was calculated following (17), and SNPs with p-value < 0.05 were considered as cis-eQTLs for the gene.

### TF-drug association tests

Hypergeometric tests were used to identify TFs whose ‘binding-change SNPs’ are enriched in drug response-associated SNPs. We used SNPs in GM12878 DNase-seq narrow peaks (28,29) as the universe. For each drug, we defined genes whose expression correlates with drug response (EC50) with a correlation p-value of 0.05 or lower as ‘drug response genes’ and eQTLs assigned to these genes (see above) as the drug response-associated SNPs. A TF’s ‘binding-change SNPs’ were defined as those with large TFBS-SNP impact score using either MOP or gkmSVM. In particular, SNPs located within the TF’s ChIP peaks and ranked among the top 300 by delta-MOP or among top 300 by delta-gkm-SVM score were called ‘binding-change SNPs’.

## RESULTS

### Selection of methods for predicting impact of SNPs on TF-DNA binding

We first sought a method to predict the impact of a SNP on TF binding (henceforth referred to as the ‘TFBS-SNP impact prediction task’), with the ultimate goal of utilizing such predictions to discover TF-phenotype relationships. This requires a sensitive method to quantify the strength of binding, since the effect of a typical SNP on a binding site is expected to be relatively modest. Several such methods have been reported in the literature (10,13,30), including some that utilize a variety of data types, such as chromatin state profiles (31) and high-resolution DNA accessibility (15,31), for prediction (10,13,30). To ensure wide applicability, we were specifically interested in a method that can predict TF binding strength from DNA sequence alone, while possibly using ChIP-seq data for the TF for model-training purposes. Existing tools for this scenario rely either on the k-mer composition of sequences (10,13,30,32) or use pre-determined motifs for the TF (33–36); recent evaluation (22) on allele-specific binding (ASB) data suggests that the k-mer based methods have a clear advantage over motif-based methods. However, the motif-based methods tested by Wagih et al. (22) use a relatively rudimentary notion of motif matching, while past work by us (26) and others (36) has contributed more sophisticated biophysical models for this purpose. We compared a representative of leading k-mer based methods (gkm-SVM (10,11)) with an advanced motif-based method to determine their relative merits in predicting TF binding strengths and their changes due to SNPs.

We first used the thermodynamics-based method called Sequence To Affinity Prediction (STAP) (26) and trained it on ChIP-seq data for a TF, thereby learning to predict the strength of TF binding (ChIP signal strength) at a putative site from its sequence and the TF’s motif. STAP scores a genomic window, e.g., a few hundred base pairs long – the typical length of a ChIP peak – for its estimated occupancy by a TF, using the latter’s pre-determined motif in position weight matrix (PWM) form. We have previously used this approach to accurately model ChIP data in D. melanogaster (26) and in mouse ESCs (37), as well as in the human cell line data sets of a recent ‘DREAM’ challenge. However, we recognized that often there are multiple motifs for the same TF in the literature or databases and it is not clear which one of them, if any, is the optimal motif to use for the modeling of binding strengths. We therefore trained separate STAP models for each available motif for a TF, and then used a support vector machine (SVM) classifier to combine the binding strength predictions of a TF at a given genomic window, made by those STAP models, into a single score (Fig 1a). We call this the ‘MOP’ (Motif-based Occupancy Prediction) score. With a means to score a window for its strength of TF binding, we were able to estimate the effect of a SNP by considering a 500 bp window centered on that SNP position, scoring two versions of the window, with the central position being set to either allele of the SNP, and computing the difference (Fig 1b). We refer to this as the ‘Delta-MOP’ score of the SNP for the TF. Note that this score is tied to the cell type from which ChIP data used in training were obtained.

**Figure 1.**
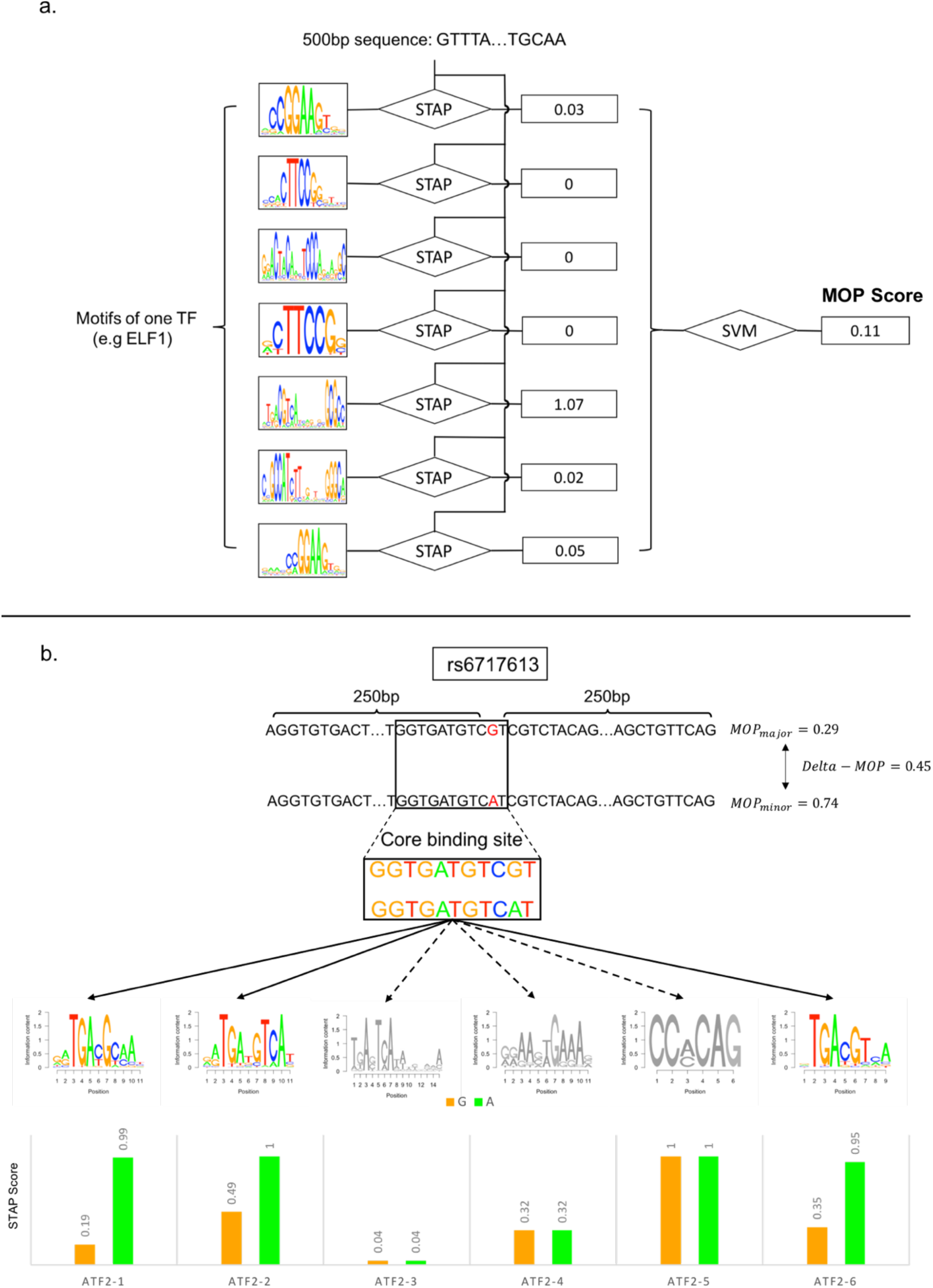
Process of scoring TFBS-SNP impact and identifying a TF’s ‘binding change SNPs’. a) We build a STAP model to predict TF binding at a DNA segment, separately for every available motif from ENCODE, FactorBook and HOCOMOCO that represents the TF. For a given sequence, each motif-specific STAP model outputs a score indicating the occupancy of the TF on the sequence. An SVM model then combines STAP scores from all motifs of the TF to compute a combined score of the TF’s binding to the sequence; this is called the ‘MOP’ score. b) ‘Delta-MOP’ score of a SNP is defined as the absolute value of the difference between the MOP scores of the major and minor allele sequences, constructed from the 501 bp sequence centered on the SNP location. In this example, SNP rs6717613 (G->A) is found to have a Delta-MOP score of 0.45 for the TF ATF2, which is the difference of MOP scores between the major and minor alleles (0.29 and 0.74 respectively). MOP scores were based on combining scores for six different ATF2 motifs (logos shown). The Delta-MOP score in this example can be qualitatively understood in terms of matches of the core binding site (top) to each of the six ATF2 motifs, whose STAP scores are shown separately for the two alleles (bottom). The core site’s match to motifs ATF2-1, ATF2-2 and ATF-6 changes in strength between the two alleles. For instance, the SNP falls on the 10th position of motif ATF2-1, which prefers an ‘A’, and the change from ‘G’ (major allele) to ‘A’ (minor allele) is interpreted as a change in strength of motif match. On the other hand, the core site does not have a strong match to ATF2-3 or ATF2-4, in either allelic form, while motif ATF2-5 overlaps the core site but not the SNP position. The Delta-MOP score combines these different pieces of information in a principled manner to compute an overall score of the impact of rs6717613 on ATF2 binding.

Figure 1b illustrates the Delta-MOP score with an example. The SNP rs6717613 (G->A) is assigned a Delta-MOP score of 0.45 for the TF ATF2, with the MOP scores of the G and A alleles being 0.29 and 0. 74 respectively. Note that six different motifs were available for this TF; for three of these ATF2 motifs, the SNP position coincides with an informative position of the motif and the two alleles define motif matches of differing strengths, while for the remaining three motifs, the two alleles present equally weak or equally strong sites. Hence, it is not clear a priori if this SNP should be considered as impacting binding strength or not, and it is instructive to have the Delta-MOP score provide an affirmative and quantitative answer.

### Evaluations of SNP impact prediction scores

We first evaluated methods for prediction of TF binding strength from sequence, since this underlies the prediction of TFBS-SNP impacts. As noted above, the newly developed MOP score, which underlies Delta-MOP, is a generalization of the motif-based STAP method (26,37,38) for predicting a TF’s binding strength. We therefore hoped to confirm that this generalization indeed improves the prediction accuracy. We were also interested in a leading k-mer based tool for predicting TF binding from sequence. We therefore considered the ‘gkm-SVM’ method, which has been demonstrated to be among the best for this purpose – on par (39) with deep learning-based methods such as DeepBind (12) and DeepSEA (13).

We trained the three methods – STAP, MOP and gkm-SVM – using the same training data set composed of 800 positive sequences (ChIP peaks of a TF) and 800 negative sequences (non-peaks), and cross-validated them on a set of 400 unseen sequences, balanced between the positive and negative classes. The negative sequences were randomly selected from the ChIP peaks of any other TF aside from the one under consideration (test TF); this is an important distinction from past benchmarks for the task (e.g., a recent ‘DREAM challenge’ *(ENCODE-DREAM in vivo Transcription Factor Binding Site Prediction Challenge*. Available from: http://dreamchallenges.org/project/encode-dream-in-vivo-transcription-factor-binding-site-prediction-challenge/), and was designed to make the evaluation more specific to the unique binding behavior of the test TF rather than more general properties of TF binding implicit within ChIP data, such as DNA accessibility. Our tests were performed for each of 37 different TFs, selected based on the availability of ChIP-seq data for a well-studied lymphoblastoid cell line (LCL), GM12878, and other relevant criteria (see Supplementary Note 1). We noted that MOP and gkm-SVM produce similar accuracy (Fig. 3a) on average across the 37 data sets (TFs), while exhibiting some level of complementarity. MOP shows a clear improvement over STAP (Fig. 3b, paired T-test p-value 0.0038), demonstrating the value of using multiple motifs when available. (Supplementary Table 1 tabulates the number of motifs available for each TF.)

We next evaluated the above methods for the TFBS-SNP impact prediction task, by asking if the SNPs with strongest effects on predicted TF binding, henceforth called ‘binding-change SNPs’, are enriched for allele-specific binding sites (ASB), defined as sites where ChIP-seq read counts are significantly different between alleles (22). The Delta-MOP score of the previous section is one way to predict binding-change SNPs, but analogous predictions can be made using STAP or gkm-SVM in place of MOP to score binding strengths of the two alleles. We refer to these as ‘Delta-STAP’ and ‘Delta-gkm-SVM’ (10,11) scores respectively. As a baseline, we also evaluated a fourth method, called ‘Delta-PWM’, which is a PWM-based scoring metric included in the evaluations by Wagih et al. (We used the “delta raw score” provided by them as this baseline.) We obtained allele-specific binding (ASB) data for 16 TFs in Lymphoblastoid cell lines from Wagih et al. (22), and tested whether the four above-mentioned methods can accurately discriminate ASB SNPs from non-ASB SNPs (see Methods). Performance was measured using the area under the receiver operating characteristic curve (AUROC, ROC curve of RUNX3 was shown in Fig. 3c) and the area under precision-recall curve (AUPRC), following (40). In AUROC comparisons (Fig. 3d, Supplementary Table 2), Delta-MOP appears to have better performance than Delta-STAP (average difference of 0.020, paired T-test p-value 0.0013) and Delta-PWM (average difference of 0.025), but not as significantly different from Delta-gkm-SVM (average difference of 0.0043). The median AUROC using Delta-MOP is 0.60, and that using Delta-gkm-SVM is 0. 58. Two of the 16 TFs – BHLHE40 and EGR1-had their ASB events predicted with AUROC of ~0.7 or greater when using Delta-MOP. These two methods exhibited a fair degree of complementarity in their performance on different TFs (Fig. 3d). In AUPRC comparisons (Fig. 3e), the performance of Delta-MOP is significantly better than Delta-PWM (average difference 0.095, paired T-test p-value 0.00072), but similar to the other two methods, with the medians of Delta-MOP, Delta-gkm-SVM, and Delta-STAP being 0.39, 0.38, and 0.36 respectively.

To summarize the evaluations reported above, we found that the motif-based method MOP and the k-mer based method gkm-SVM are equally good predictors of binding strength as well as of allele-specific binding events, with noticeable degree of complementarity to each other, while MOP shows clear improvements over the two other motif-based methods evaluated. We therefore selected Delta-MOP and Delta-gkm-SVM to predict TFBS-SNP impact for the next steps of analysis. It was instructive to find that a motif-based approach (Delta-MOP) is competitive with, and for some TFs better than, the k-mer based Delta-gkm-SVM method (see Discussion).

### Discovery of TFs regulating individual variation in cytotoxic drug response

To discover TFs associated with phenotypic variation, we adopted a statistical approach illustrated in Fig. 2. At its heart is a Hypergeometric test of the overlap between two sets of SNPs, outlined below.

**Figure 2.**
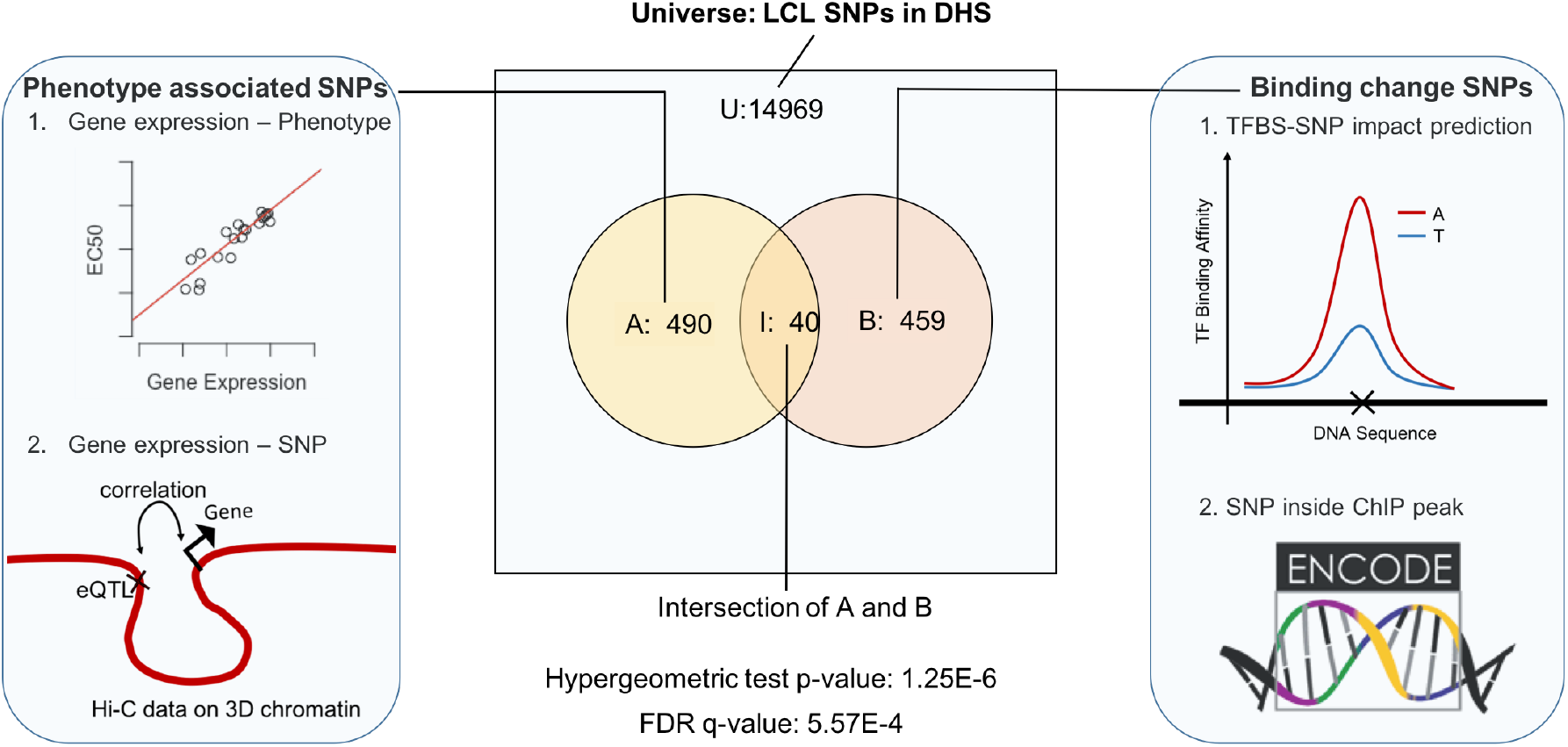
Process of identifying TFs regulating phenotypic variation. A hypergeometric test is used to test the overlap between a TF’s ‘binding change SNPs’, based on presence within ChIP peaks from ENCODE and high Delta-MOP score, and ‘phenotype-associated SNPs’, i.e., eQTLs of genes whose expression correlates with phenotype, located within cis-regulatory regions of the gene identified by Hi-C data. A TF is considered significant to the phenotype if the FDR q-value is below 0.05.

TF-phenotype association test:

a. We consider the collection of all SNPs that are located within accessible DNA in the cell type of interest; this is the ‘universe’ set for the test. (Also, see Discussion about this choice.)
b. We define a subset of SNPs that are likely to impact phenotypic variation through a cis-regulatory effect on a gene’s expression; we refer to this as the ‘phenotype-associated’ SNPs. Specifically, we identify phenotype-associated genes based on significant association between gene expression and the phenotype, and then determine significant eQTL SNPs in the regulatory regions (explained below and in Methods) of those genes.
c. We separately define a subset of SNPs that are likely to affect a particular TF’s binding strength, i. e., the ‘binding-change’ SNPs. Although introduced above, these are now redefined as the SNPs with the greatest Delta-MOP or Delta-gkm-SVM score for that TF, among those located within the TF’s ChIP peaks for the cell type (see Methods).
d. A Hypergeometric test is used to test the overlap between phenotype-associated SNPs and binding-change SNPs; a significant overlap is considered as evidence for the TF’s role in regulating phenotypic variation.

We note that the above test, conducted at the level of SNPs, is conceptually similar to that in Hanson et al. (16), with several key differences, the most prominent being our use of TFBS-SNP impact prediction scores as an additional criterion for designating binding-change SNPs. Hanson et al., in contrast, considered all SNPs within the TF’s ChIP peaks as binding-change SNPs. Other important differences are that Hanson et al. performed the statistical test at the gene level and did not use DNA-accessibility or enhancer-promoter interaction data.

We used the TF-phenotype association test procedure on a data set of 284 lymphoblastoid cell lines (LCLs) that have previously been assayed for their cytotoxic response (EC50) to each of 24 different treatments, mostly cancer drugs (17). Gene expression and genotype data are also available for these LCLs. We used ENCODE (41–43) ChIP-seq data for 37 TFs in the lymphoblastoid cell line GM12878 (see Methods), along with the above-mentioned genotype data to identify binding-change SNPs, using Delta-MOP and Delta-gkm-SVM for TFBS-SNP impact prediction. We also repeated the analysis using only one or the other of these methods (see Supplementary Tables 3,4). To identify phenotype-associated SNPs, we considered genes whose expression levels correlated significantly with EC50 values (of a specific drug) across the panel of LCLs, and used Hi-C data (21) from the GM12878 cell line in step (b) of the above procedure. Here, we defined the regulatory region of a gene to include the chromatin interaction domain to which the gene belongs (see Methods), as well as more distal segments predicted to interact with the gene via chromatin ‘loops’. (21)

### Assessment of predicted TF-drug associations

A total of 888 TF-drug pairs (24 drugs x 37 TFs) were evaluated; we report in Table 1 all 38 pairs significant at False Discovery Rate (FDR) of 5% (nominal p-value < 0.0021). (The full results are in Supplementary Table 5.) We also performed a variant of the above enrichment tests where TFBS-SNP impact prediction was not used; instead a size-matched set of randomly selected SNPs within ChIP peaks (of the test TF) were chosen for consideration as binding-change SNPs, as was done by Hanson et al. (16). (We used a size-matched random subset of within-peak SNPs, rather than all such SNPs, so that enrichment levels can be compared.) We repeated this ‘randomized control’ test 100 times and noted how frequently each significant pair in the original analysis had a stronger p-value in these randomized controls, reported in Table 1 (column ‘Impact predictor utility p-score’). We note that 21 of the 38 reported pairs have only ≤ 10% chance of being discovered when not using TFBS-SNP impact prediction scores, thereby underscoring the value of such predictions in the procedure. This comparison establishes that impact prediction scores can indeed help identify novel statistical associations, though a rigorous assessment of the sensitivity-precision tradeoff due to their use is not attempted here. In another control experiment, we assigned to each TF a random set of SNPs (size-matched with the binding-change SNP sets above) from the universe of all SNPs within accessible regions and tested all 888 TF-drug pairs. We discovered that, on average, across 100 such randomized control tests, only 1. 27 pairs (about 0.14% of the 888 tested) were significant at a nominal p-value of 0.0021, the criterion used above for reporting (Table 1), providing further statistical evidence for the low proportion of false positives in our report.

**Table 1.** Significant TF-Drug associations. 38 TF-Drug pairs were discovered as significant at False Discovery Rate (FDR) of 5% (nominal p-value < 0.0021). P-value of the hypergeometric tests are shown in the third column. The fourth column (‘Impact predictor utility p-score’) shows an empirical p-value for each association, computed by repeating the hypergeometric test using a size-matched random subset of SNPs within ChIP peaks (rather than SNPs with greatest TFBS-SNP impact scores) 100 times and counting how frequently the test p-value in these random controls is smaller than that observed in the original test for that TF-Drug pair.

| TF | Drug | p-value | Impact predictor utility p-score |
| --- | --- | --- | --- |
| SIN3A | CDDP | 3.04E-06 | 0.2 |
| ELF1 | RAPAMYCIN | 3.11E-06 | 0 |
| ELF1 | EPIRUBICIN | 4.92E-06 | 0.06 |
| MAX | CDDP | 8.74E-06 | 0.02 |
| MAX | 6TG | 1.88E-05 | 0.05 |
| SP1 | CARBOPLATIN | 1.95E-05 | 0 |
| ELF1 | DOXORUBICIN | 2.44E-05 | 0.08 |
| NFYB | 6TG | 2.94E-05 | 0.01 |
| NFYB | 6MP | 7.45E-05 | 0.08 |
| ELF1 | CARBOPLATIN | 7.84E-05 | 0.04 |
| SIN3A | MPA | 8.22E-05 | 0.06 |
| MXI1 | CDDP | 1.18E-04 | 0.12 |
| PML | EPIRUBICIN | 1.89E-04 | 0.06 |
| SIN3A | 6TG | 1.99E-04 | 0.66 |
| SP1 | CDDP | 2.07E-04 | 0.03 |
| YY1 | CDDP | 2.15E-04 | 0 |
| ELF1 | CDDP | 2.41E-04 | 0.11 |
| REST | 6MP | 2.98E-04 | 0.25 |
| REST | RAPAMYCIN | 3.07E-04 | 0.6 |
| SP1 | EPIRUBICIN | 4.66E-04 | 0.06 |
| NFYB | EPIRUBICIN | 4.88E-04 | 0.2 |
| BHLHE40 | TMZ | 5.23E-04 | 0.02 |
| PBX3 | EPIRUBICIN | 6.00E-04 | 0.01 |
| MXI1 | 6MP | 6.29E-04 | 0.04 |
| MAX | 6MP | 6.88E-04 | 0.22 |
| SP1 | OXALIPLATIN | 6.89E-04 | 0.01 |
| PML | DOXORUBICIN | 7.08E-04 | 0.14 |
| MAX | EPIRUBICIN | 8.25E-04 | 0.24 |
| NFYB | DOXORUBICIN | 8.28E-04 | 0.23 |
| REST | CARBOPLATIN | 8.46E-04 | 0.22 |
| ATF2 | FLUDARABINE | 1.04E-03 | 0 |
| BHLHE40 | EPIRUBICIN | 1.21E-03 | 0.01 |
| MXI1 | CARBOPLATIN | 1.60E-03 | 0.22 |
| SP1 | DOXORUBICIN | 1.61E-03 | 0.11 |
| ATF2 | EVEROLIMUS | 1.77E-03 | 0 |
| PAX5 | HYPOXIA | 1.86E-03 | 0.03 |
| SIN3A | MTX | 1.93E-03 | 0.23 |
| PBX3 | DOXORUBICIN | 2.05E-03 | 0.01 |

While we showed above that the use of TFBS-SNP impact scores can help predict TF-drug associations that might otherwise not rise above statistical significance, we also needed to convince ourselves that the discovered statistical associations are likely to be biologically true. In the absence of any systematic benchmarks of causal relationships between TFs and drug response, we had to rely on extensive but ad hoc survey of the literature for supporting evidence, following guidelines established in (44). Out of the 38 significant TF-drug pairs of Table 1, eight were found to have ‘direct’ supporting evidence (Table 2). For seven of these 8 cases, knock-down of the TF has been shown to lead to a significant difference in sensitivity. In one case – the pair ELF1-CDDP – we found published evidence that DNA-bound ELF1 increases CDDP-induced DNA damage at the bound locations, thereby directly and mechanistically implicating the TF’s regulatory activity in response to the drug. Notably, three of the top seven significant pairs (based on p-value) have such direct confirming evidence, and these three pairs would have not have been discovered if not using TFBS-SNP impact scores (Impact predictor utility p-score <= 0.1, Table 1). Among the eight pairs with direct evidence, only two (ELF1-CDDP and PML-Doxorubicin) would have reasonable chance of being discovered without use of TFBS-SNP impact information (Impact predictor utility p-scores of 0.11 and 0.14 respectively).

**Table 2.**
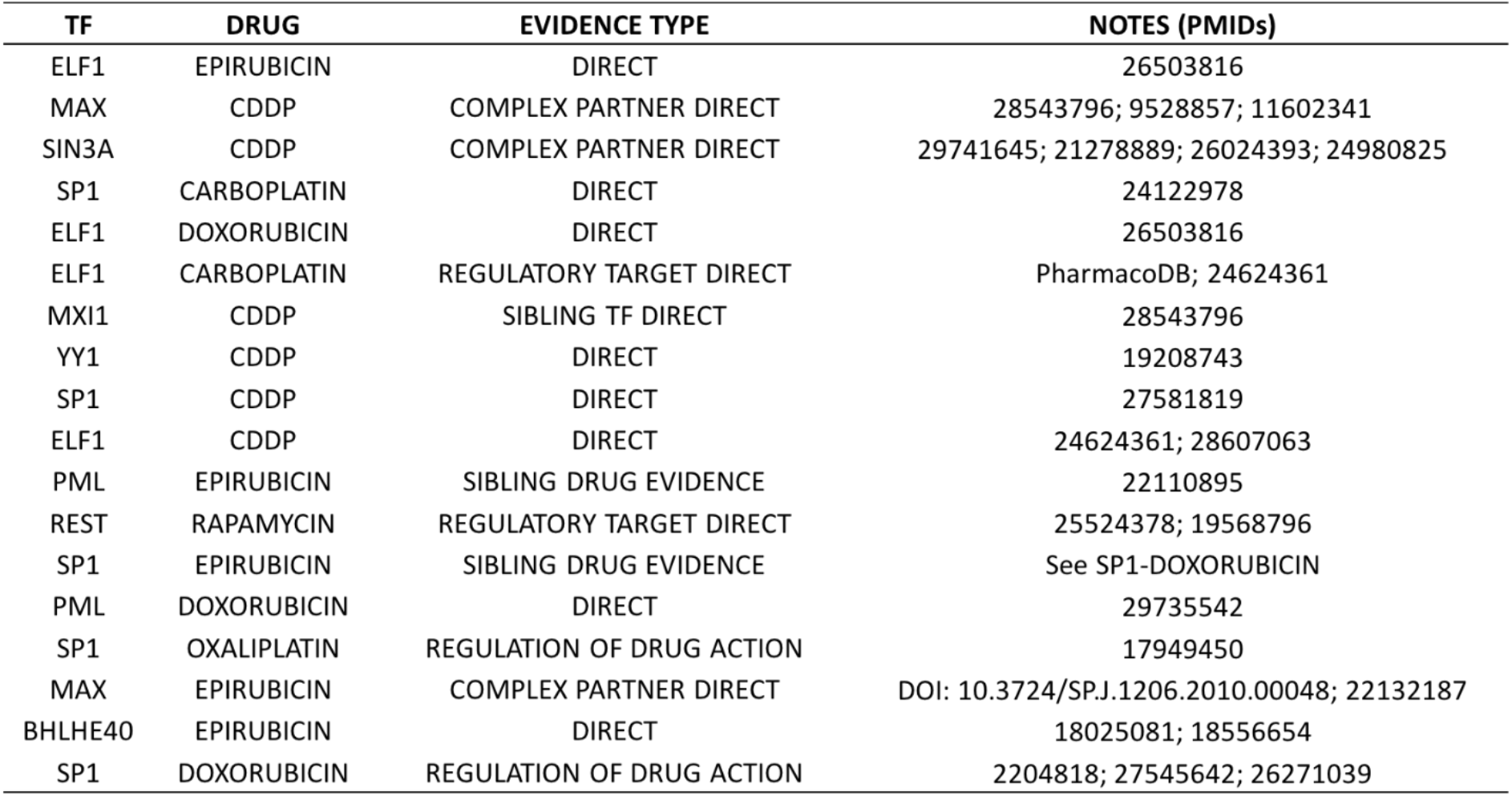
TF-Drug pairs with supporting evidence. This table lists the 18 TF-Drug pairs (among the 38 pairs shown in Table 1) that have supporting literature evidence. We defined four different evidence types based on the type of evidence, as explained in text.

We found six additional pairs to have strongly suggestive evidence of a biological relationship. This includes cases where the TF is a demonstrated regulatory mechanism of the drug’s action (evidence code ‘Regulation of Drug Action’ in Table 2), is a known regulator of the drug’s target protein or pathway (‘Regulatory target direct’ in Table 2), or plays a role in sensitivity to a closely related drug (‘Sibling Drug Evidence in Table 2); see Supplementary Note 3 for details. As an example of ‘Regulation of Drug Action’, SP1-mediated trans-activation of survivin has been shown to reduce Doxorubicin sensitivity (45), supporting the pair SP1-Doxorubicin. An instance of ‘Regulatory Target Direct’ evidence is provided by the pair REST-Rapamycin: REST is known to exert regulatory control over the ‘mTOR’ signaling pathway (46) and this pathway (mTOR = ‘mammalian Target Of Rapamycin’) is the canonical target of the drug Rapamycin (47). An example of the evidence code ‘Sibling Drug Evidence is the pair PML-Epirubicin, supported by direct evidence for the role of TF PML in response to the drug Doxorubicin, which is closely related to Epirubicin (48) and is expected to have very similar mechanisms to the latter.

In three additional cases, we found direct evidence in favor of a physical interaction partner of the implicated TF having an effect on the drug (evidence code ‘Complex Partner Direct’). For instance, while the reported pair MAX-CDDP does not have direct evidence, the ‘Max Dimerization Protein 1’ (MXD1), a member of the Myc-Max-Mxd family, is a well-known dimerization partner of MAX (49), and has been shown to induce CDDP (cisplatin) resistance in hypoxic U 2OS and MG 63 cells (50). As another example, SIN3A is part of the SIN3A-HDAC complex that is associated with diseases including cancer (51), and HDAC inhibitors are known to potentiate CDDP activity (52–54), thus providing moderate but indirect evidence in favor of the reported pair SIN3A-HDAC. An additional example of similar indirect evidence was found for the pair MXI1-CDDP: MXI1, also known as MXD2, is a member of the MXD1 family, and a closely related member of this family – MXD1 – has been shown, via knockdown assays, to affect CDDP resistance in hypoxic conditions via repression of PTEN (50).

Thus, we were able to retrieve direct or indirect evidence in support of 18 of the 38 reported TF-drug pairs of Table 1. Eleven of these 18 are significant only when using the TFBS-SNP impact prediction scores (Impact predictor utility p-score <= 0.1), making the case for the added value of these scores in cis-regulatory analysis leading to mechanisms of phenotypic variation. Some TFs and drug families were clearly overrepresented in the predictions of Table 1. For instance, ELF1 was predicted to be associated with five drugs, with direct evidence for three of these associations, and indirect evidence for a fourth. The TF SP1 was also associated with five drugs, two of which are supported by direct evidence and the remaining three by indirect evidence. The two anthracyclines included in the tests – Doxorubicin and Epirubicin – accounted for 12 of the 38 predicted pairs, with four supported by direct and four by indirect evidence. The platinum therapy drug CDDP (cisplatin) was found associated with six TFs, and all these associations were supported by the literature (three by direct and three by indirect evidence).

### Regulatory mechanisms underlying variation in doxorubicin response in LCLs

Our statistical procedure not only identifies TFs likely to regulate a drug’s cytotoxic response, it also provides us the opportunity to probe more deeply into the regulatory interactions mediating such a TF’s influence. Each TF-drug pair is reported based on a statistically significant overlap between the drug-response SNPs and the binding change SNPs. Thus, the SNPs in this ‘intersection set’ represent the confluence of four pieces of evidence: they are located within ChIP peaks of a TF, have evidence suggesting impact on TF binding site strength, are statistically correlated with the expression of a cis-linked gene (via chromatin interaction), and the gene’s expression, in turn, is correlated with drug response levels. A fifth important piece of evidence is that the TF is likely to be a regulator of the phenotype (in light of the significant p-value), especially if the association is also supported by prior literature evidence. Thus, we considered the SNPs in the above-mentioned ‘intersection set’ as especially informative, and examined the genes linked to them for further evidence of phenotype-relevance.

We report our findings for the pair ELF1-Doxorubicin, a statistically significant association that is also supported by direct experimental evidence in the literature. (It was also one of the TF-drug pairs that did not rise to significance when repeating the TF-phenotype association test without using TFBS-SNP impact prediction.) The SNPs in the intersection set for this pair were linked to 39 unique genes (Supplementary Table 6), which are putative mediators of ELF1 influence on drug response. We first reconstructed a skeleton ‘pathway’ of major known events in Doxorubicin-induced cell death (Figure 4, rectangles, solid black edges, and ovals placed on these edges), based on the literature (55). This mainly involves DNA damage by topoisomerase II (TopII) inhibition and generation of reactive oxygen species (ROS) through a redox reaction involving the free radical semiquinone (56). Genes known to be important for DNA damage-induced cell death include TP53, ATM, BCL2, BCL2L13, BAX, BAK1 and P21 among others (57,58), while those involved in transduction of ROS signals include RAS, RAF, MEK, ERK, and P38, among others (59). Both routes of Doxorubicin-response involve CASP3 and CASP9 as an apoptotic mechanism (60).

We sought to relate each of the 39 genes identified above to this pathway via known regulatory interactions. We were successful in finding such relationships for 15 of the 39 genes (shown in Figure 4 as green ovals and dashed edges). (See Supplementary Note 4 for details). For instance, the genes ATM and BAK1 were identified as potential mediators in our hypergeometric test, and are in the skeleton pathway constructed above. ATM is part of the ATM/TP53 pathway and plays an important role in the activation of TP53 (55), while BAK1, a member of the BCL2 protein family, is activated by TP53 and is known to induce apoptosis (55). The microRNA miR-6734 is an identified ELF1-Doxorubicin mediator, and is known to up-regulate the expression of P21, which is a TP53-inducible apoptosis inhibitor in our skeleton pathway (61). Another gene, NEURL4 which is also identified as a mediator, has been shown to be a regulator of TP53 activity. Another potential mediator, B4GALT2, has been identified as a regulatory target of TP53 and plays a role in DNA damage-induced apoptosis. Interestingly, binding sites of ELF1 have been identified in the cis-regulatory region of B4GALT2, further supporting its predicted role as a ELF1 mediator (62). As revealed by these and additional examples shown in Figure 4 (also Supplementary Note 4), our procedure can not only identify major TFs regulating phenotypic variation but also several of the genes mediating its influence, via a subset of SNPs that have multiple lines of supporting evidence.

## DISCUSSION

We have examined the challenging problem of non-coding variant interpretation, using a motif-based computational method to predict TF-DNA binding impact, followed by assessment of putative high-impact variants for potential links to phenotypic variation among individuals. Our major contribution is in showing that use of binding impact prediction can help identify regulatory mechanisms (identities of TFs) relevant to phenotypic variation, that would be missed if relying only on the location of variants inside TF binding sites. In doing so, we have begun to bridge the actively researched field of TFBS-SNP impact prediction (15) with genotype-phenotype association studies (1,63), especially those that simultaneously examine genotype, gene expression and phenotype data from a cohort. It is common for authors of cis-regulatory impact prediction tools to test if their predictions are enriched in GWAS and other disease-related SNPs (19), in efforts to build exactly such a bridge. Our approach takes the idea a step forward by making it the goal rather than a means to validate predictions. That is, rather than stop at predictions of high impact SNPs and use GWAS enrichments as a sign of significant predictive performance, we solve a problem – identify TF-phenotype associations – that crucially depends on the second step. To be clear, our enrichment tests are not against GWAS SNPs, rather against eQTLs for phenotype-correlated genes; this seems more in line with the expectation that SNPs with high predicted impact on TF binding should link to phenotype by causing expression variation. Moreover, since the impact predictor (Delta-MOP or Delta-SVM) was trained on LCLs, and cell line-specific predictors are indeed the norm today (10), linking them directly to GWAS SNPs for a particular disease is premature and has to cross the generalization barrier from cell lines to tissues where disease-related dysregulation happens.

The hypothesis testing approach to TF-phenotype associations was adapted from our previous work (16), with the key difference being the use of TFBS-SNP impact prediction as an essential part of the approach; Hanson et al. assumed that a SNP located inside a ChIP peak is evidence for the TF’s potential regulatory effect on the nearby gene’s expression. We add the criterion of high TFBS impact (as predicted by Delta-MOP or Delta-gkmSVM) as a requirement for SNPs mediating TF regulatory control, thus making the evidence more reliable and also opening up a link to the rapidly maturing field of cis-regulatory impact prediction. Our direct comparisons between results (TF-drug associations) obtained with and without use of Delta-MOP scores (Table 1) are among the first direct statistical findings of the value of TFBS-SNP impact prediction for mechanistic studies of phenotypic variation, especially on genome-wide scales. More ‘localized’ applications, such as prioritization of a small number of candidate SNPs, are already being reported in the field (5). We also note that the basic methodology of our work can serve as a practical way to assess the value of new approaches to SNP scoring and prioritization, since the results of this methodology are testable findings at the TF and gene level (their roles in phenotype, as illustrated in Table 2 and Figure 4): there is more literature evidence to compare against at these levels than there is evidence for SNP function and mechanism. A technical note related to the hypothesis testing framework (Hypergeometric test) is that the universe comprises all SNPs within accessible regions of DNA, rather than all SNPs. This choice was intentional, since both subsets tested for overlap – the phenotype-associated SNPs and the binding-change SNPs – are expected to be highly enriched in accessible regions, the former because they are eQTLs and the latter because they are within TF ChIP peaks. Using accessibility as a required criterion for all SNPs in the universe thus factors out the effect of accessibility in the analysis, focusing on the TF’s regulatory role more directly.

We also developed a new computational method, called MOP, for predicting TF binding strength, which underlies the Delta-MOP score for TFBS-SNP impact prediction. There are several tools available today for predicting TF binding from sequence. We chose to build MOP, based on a thermodynamics-based method called STAP (26), partly because we have extensive experience with the latter. But there were additional motivations for this choice. For example, due to its biophysics-inspired formulation and a single free parameter (that reflects TF concentrations), the STAP method offers a more interpretable scoring compared to highly parameterized k-mer based models (10,11) and deep learning models (12,13). For the same reason, it is more amenable to future incorporation of additional mechanisms specific to a TF’s binding strength, e.g., frequent cooperative binding with a secondary TF (64–66). On a different note, we have been intrigued by recent reports that k-mer based models are clearly superior to motif-based models, for TFBS impact prediction. However, the most comprehensive such evaluation in our knowledge – that by Wagih et al. (22) – used a fairly basic motif-based approach to compare against, and we sought to confirm the claim ourselves. We therefore performed systematic comparisons, for prediction of TF binding (Figure 3 a-b)as well as binding impact (Figure 3 d-e), between MOP – a multi-motif extension of STAP – and gkm-SVM – a popular, mature and easy-to-use tool that performed as well as any other evaluated in (22). Our tests suggest that the motif-based MOP and the k-mer based gkm-SVM have very similar performance, and thus prompt a re-examination of the merits and flaws of motif-based methods. For our purposes, the observed complementarity between the methods (Figures 3) was a promising finding that suggested that we use both of them in our downstream, phenotype-related analyses (Table 2). Finally, we note that the new capability of MOP compared to STAP is the automated handling of multiple motifs of the same TF. This solves a practical problem with many motif-based methods, since the same TF often has several motifs identified through different technologies and algorithms (67–69). We also extended the STAP method to predict binding using motifs for other TFs, but did not observe significant advantages and leave further development of the idea for future work (data not shown).

**Figure 3.**
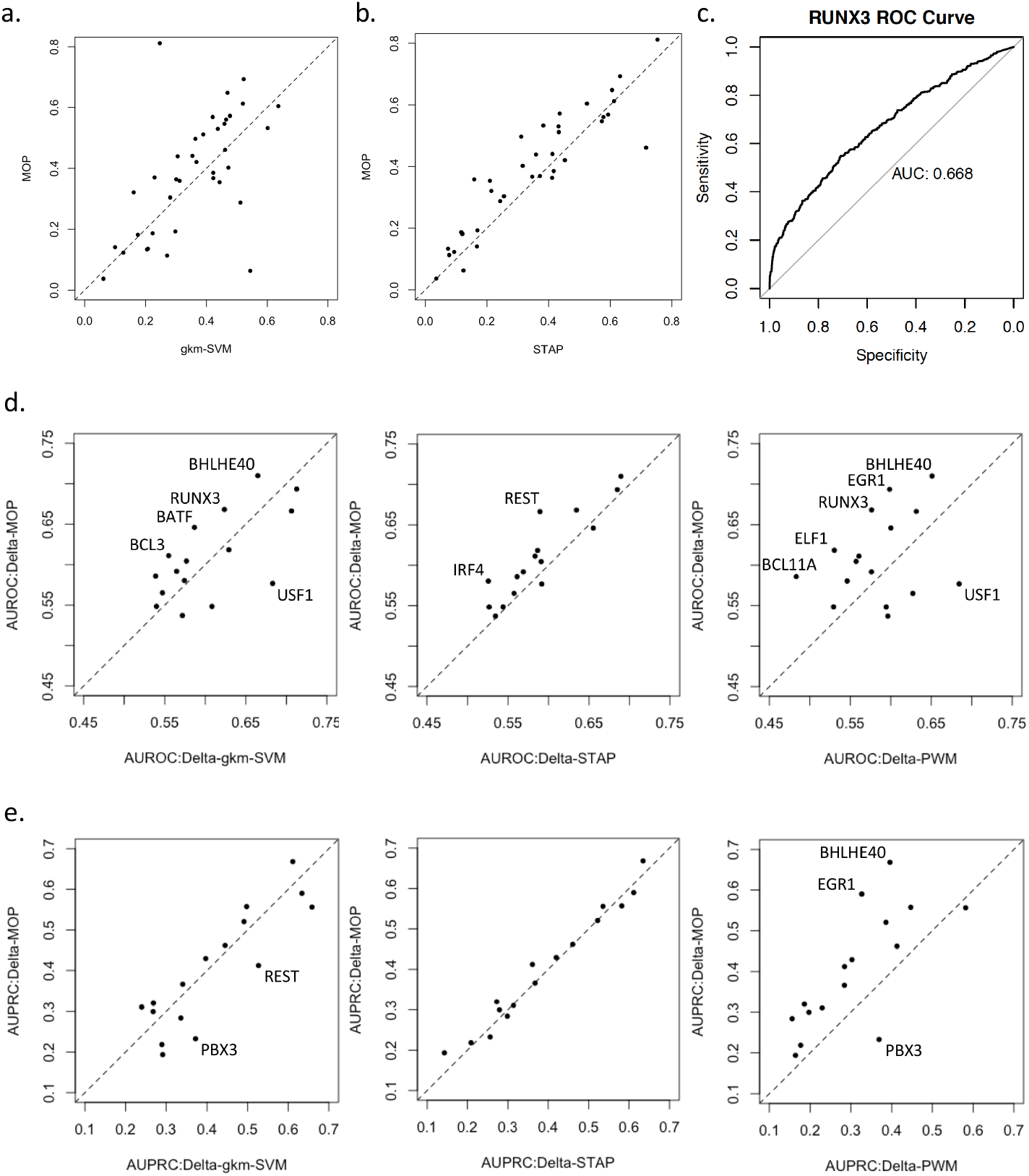
(a-b) Comparison of three TF binding predictors. We compared MOP with STAP and gkmSVM. The performance of each model is measure by the Pearson Correlation Coefficient (CC) between ChIP score and predicted binding score on a test set of 400 sequences that are not used in model training. Performance evaluation is performed for each of 37 data sets (for different TFs). a) MOP performs as well or better than STAP (using the best motif when multiple motifs are available) for 26 of the 37 data sets, with their average CC being 0.39 and 0.36 respectively. b) MOP performs as well or better than gkmSVM for 21 of 37 TF data sets examined, with average CC of the two methods being 0.39 and 0.37 respectively. (c-e) Evaluation of TFBS-SNP impact prediction methods. Four different methods of binding change prediction (Delta-MOP, Delta-gkm-SVM, Delta-STAP and Delta-PWM) were evaluated for their ability to predict allele-specific binding (ASB) events from non-ASB events, for each of 16 data sets based on ChIP-seq data for different TFs. Performance was measured using AUROC as well as AUPRC. ROC curve of RUNX3 using ‘Delta-MOP’ as impact predictor is shown in (c). The last two rows show pairwise comparison of Delta-MOP and each of the other three methods based on AUROC (d) and AUPRC (e) achieved by the methods on the same data set.

**Figure 4.**
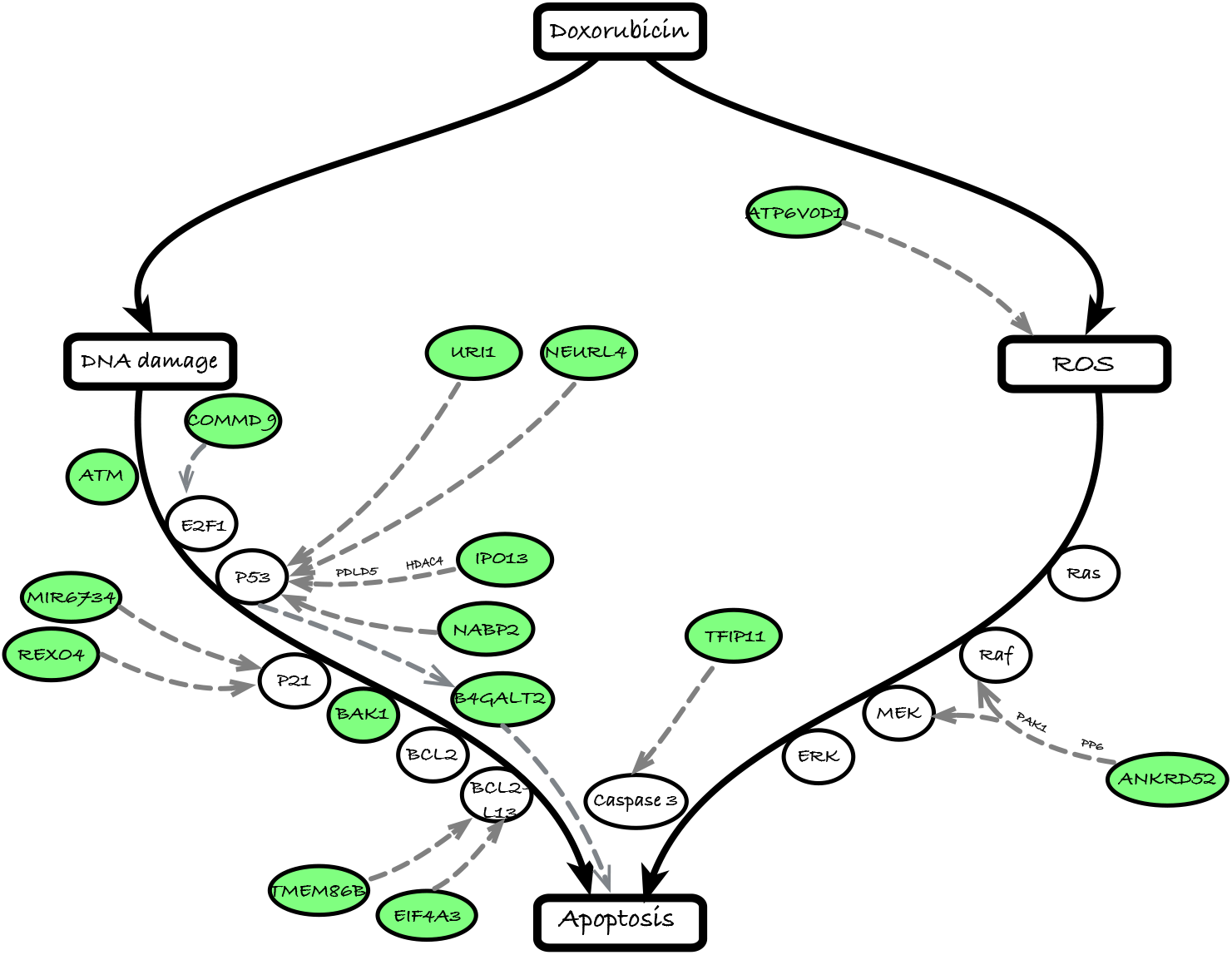
Predicted mechanisms of ELF1-regulation of Doxorubicin-induced apoptosis. Black solid arrows show the skeleton of two major pathways to Doxorubicin-induced apoptosis, viz., those mediated by DNA damage and Reactive Oxygen Species (ROS) respectively. Genes directly involved in these pathways are shown as ovals placed on the arrows. Green ovals represent drug response-associated genes that are predicted to be regulated by ELF1 and have been previously shown to have regulatory function on a pathway gene. Such regulatory evidence, presented in previous literature, is represented by grey dashed arrows connecting ELF1-regulated DRGs to pathway genes.

A note is in order regarding our overall scheme for training and testing binding predictors such as MOP and gkm-SVM: contrary to the more common practice of discriminating between ChIP peaks of a TF and random segments of the genome (70), we train (and test) models using ChIP peaks of the TF as the positive class and ChIP peaks of other TFs as the negative class. We believe this approach to testing better reflects the ability of a model to capture the test TF’s binding features in the sequence, as opposed to more general factors such as DNA accessibility that influence TF-DNA binding (71). Moreover, for highly parameterized models such as gkm-SVM, this manner of training likely results in models being better trained for prediction of a SNP’s impact on the test TF’s binding rather than TF binding in general. This is important for the downstream application (identification of phenotype-related TFs) of TFBS-SNP impact predictions in our work. Another methodological direction that we explored but do not report on is the consideration of gene regulatory networks reconstructed (using GENIE3 (72)) from expression data on suitable cell lines (ENCODE project. https://www.encodeproject.org/) in defining the ChIP peaks for training and testing binding predictors. This idea, proposed by Svetlichnyy et al. (33), was discarded as limiting the peaks to those located near putative gene targets of a TF led to far too few peaks for successful training.

The evaluation of Delta-MOP and other TFBS-SNP impact prediction methods using allele-specific binding (ASB) data is the most direct strategy for such evaluation available today, and more direct than, for example, enrichment with disease SNPs or eQTLs. The results leave us with a sense of measured optimism: while significant accuracy values for XX of the YY tested cases is promising, there is clearly much work cut out for the future. We refer the reader to an excellent review by Bart et al. (73), who point out several challenges that need to be overcome in accurately predicting TFBS binding impact. Furthermore, even if the impact of a non-coding variant on TF binding is accurately predicted, it does not equate to a regulatory impact on the gene, and the gap between TF-DNA binding and TF-gene regulation remains to be bridged.

In earlier stages of the work, we performed preliminary evaluations of TFBS-SNP impact predictors with eQTL data rather than ASB data that was our final choice (Figure 3). We used the collection of all eQTL SNPs located within regulatory regions of their target genes as an unbiased, albeit noisy, estimate of regulatory SNPs, and tested their enrichment in predicted binding-change SNPs. We repeated the enrichment tests using each of the three methods – Delta-MOP, Delta-STAP and Delta-SVM – for prediction of binding-change SNPs. As a baseline, we also tested their enrichment in a size-matched random subset of SNPs within ChIP peaks of that TF. To our initial surprise, we found (Supplementary Note 2) that the enrichments on the whole (i.e., over all 37 TFs evaluated) were similar for the binding-change SNPs and the randomly selected (within-peak) SNPs; this was the case regardless of the method used for predicting binding change. In hindsight however, we noted that this evaluation was flawed, despite its initial appeal. In particular, we recognized that the ‘ground truth’ of regulatory SNPs used in the test – the set of cis-eQTLs – likely reflects the action of multiple TFs, while the binding-change SNPs predicted by the three methods are TF-specific.

In conclusion, this work has taken the first steps towards demonstrating the value of TFBS-SNP impact prediction in the discovery of regulatory mechanisms underlying phenotypic variation. At the same time, the performance of the impact predictors leaves much room for improvement, and future advances in this active area of research will be greatly beneficial to the reconstruction of phenotype-associated gene regulatory networks.

## AVAILABILITY

MOP, motif-based occupancy prediction is available in the GitHub repository (https://github.com/UIUCSinhaLab/MOP)

## SUPPLEMENTARY DATA

Supplementary Data are available at NAR online.

## Supporting information

Supplementary File 1: TF Binding Motifs

## ACKNOWLEDGEMENT

The authors wish to thank Keith Stewart and Neal Cohen, as well as the Mayo Clinic Center for Individualized Medicine, the Interdisciplinary Health Sciences Institute, the Todd and Karen Wanek Program for Hypoplastic Left Heart Syndrome, and the Mayo Clinic & Illinois Alliance for Technology-Based Healthcare.

## Declaration

None of the authors have any competing interests.

