## Supplementary for "Mechanistic interpretation of non-coding variants helps discover transcriptional regulators of drug response"

**Supplementary Material**: The file contains supplementary text, figures and tables in the following order.


1. **Supplementary Note 1**: ChIP-seq data pre-processing
2. **Supplementary Note 2**: eQTL evaluation
3. **Supplementary Note 3**: Literature Validation of Significant TF-Drug pairs
4. **Supplementary Note 4**: Literature Validation of Mediator Genes
5. **Supplementary Figure 1**: ASB evaluation with entire data set (genotyped and imputed SNPs)
6. **Supplementary Table 1**: Number of Motifs for each TF
7. **Supplementary Table 2**: AUROC and AUPRC Comparisons
8. **Supplementary Table 3**: TF-Drug Analysis using Delta-MOP
9. **Supplementary Table 4**: TF-Drug Analysis using Delta-gkm-SVM
10. **Supplementary Table 5**: Full Results of TF-Drug Analysis
11. **Supplementary Table 6**: Mediator Genes in ELF1-Doxorubicin Analysis
12. **Supplementary File 1**: Transcription factor binding motifs
13. **References**

**Supplementary Note 1**: Here we describe how we pre-process ChIP-seq data for training purposes.

We used clustered ChIP (version 3) track from ENCODE UCSC as experimental evidence of TF binding. Signal scores, used as measurement of experimental TF binding strength, are collected from ChIP-seq signal files from ENCODE project. We selected 1000 TFBS for each TF as positive sequences using following steps:

1. TF ChIP peaks of GM12878 cell line from TFBS clusters were used.
2. A TF's peak in the TFBS clusters is defined as a High Occupancy Target Region (HOT Region) if there are 6 or more TFs having TFBS overlapping over half of the peak.
3. For each TF, we defined a subset of ChIP peaks from TFBS clusters as function peaks with two criterions. 1) overlap with a protein coding gene's 50kb upstream regulatory region; 2) not a HOT region.
4. We selected the top 1500 peaks with the largest ChIP score for each TF.
5. For each selected peak, signal scores for each base pair in the window were obtained from ChIP-seq signal file. The maximum signal score was used as a measurement of TF binding strength at this window. The window was then modified to be a 501bp window around the position with the maximum score (250bp upstream and 250 bp downstream). Top 1000 peaks with the largest signal score were selected to be positive windows where the corresponding TF binds.
6. Negative windows were defined as open chromosome region that are able to be bound by other TFs and show no evidence of binding of the target TF. Negative windows for $TF_i$ were selected as follows: 1) Take a union of all positive windows of all $TF_j$ $(j \neq i)$. 2) Remove windows with signal scores greater than the mean of the 1000 positive window signal scores of $TF_i$. 3) Randomly select 1000 windows from this union as negative windows of $TF_i$.

ENCODE clustered ChIP track contains ChIP-seq data for 62 TFs for GM12878. After the pre-processing steps mentioned above, the number of TFs having enough data decreased to 37. A dataset with sequences and signal scores of these 2000 windows were obtained for each TF and used for training and testing.

**Supplementary Note 2**: eQTL evaluation

Besides ASB evaluation, we also evaluated TFBS-SNP impact prediction of four methods by asking if the 'binding-change SNPs' are enriched for eQTLs using hypergeometric tests. We evaluated 'binding-change SNPs' predicted by four methods: Delta-MOP, Delta-gkmSVM, Delta-STAP and Delta-LLR and compared them with a set of same number of randomly selected SNPs that covered by the TF's ChIP peak. The scatter plots show pairwise comparison of each method (Delta-MOP, Delta-gkm-SVM, Delta-STAP and Delta-LLR) and its corresponding random test. For all four methods, the overall enrichment were similar no matter the TF binding predictors were used or not.

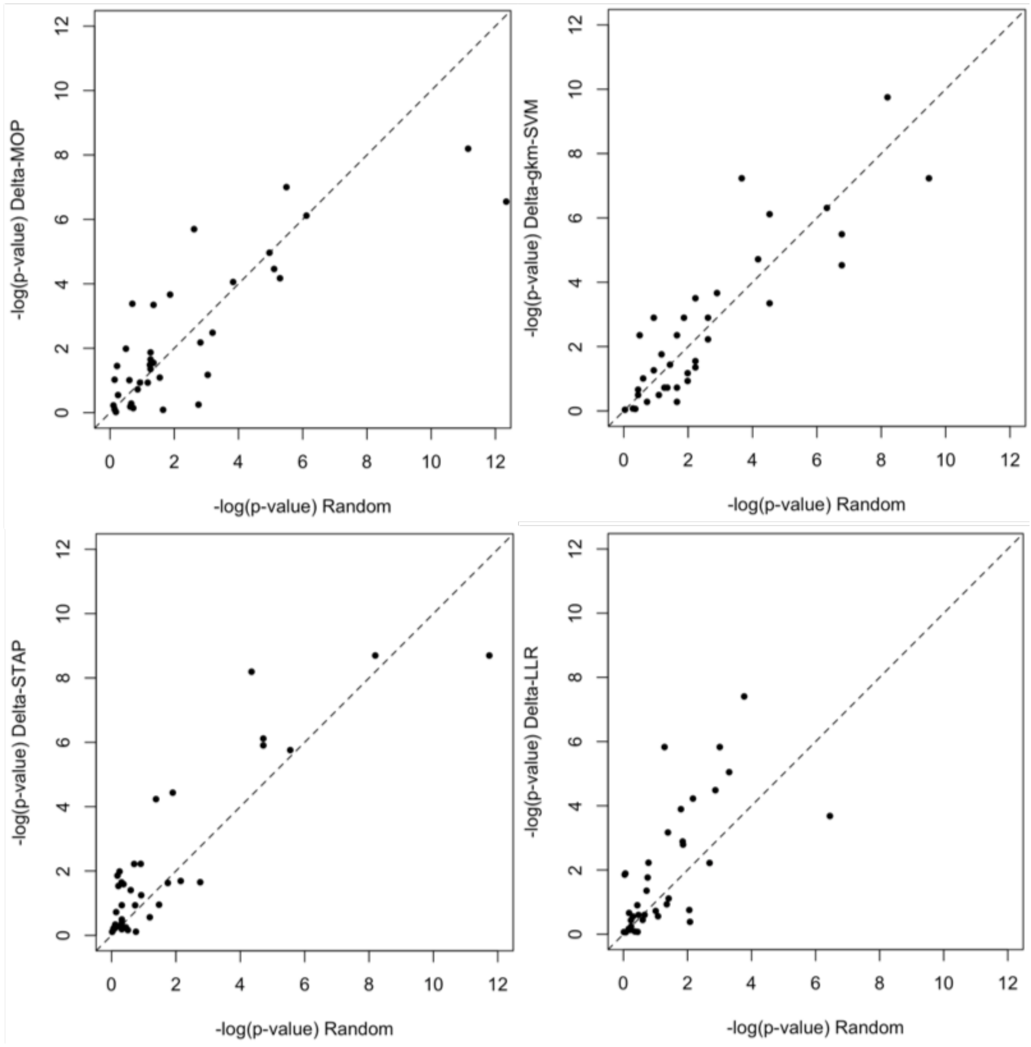

**Supplementary Note 3**: Here we list and discuss the literature evidence found for the 38 significant TF-Drug pairs identified in the TF-Drug analysis. For each TF-Drug pair, we provide an evidence code and a brief summary of the evidence. We considered five types of evidence code.

1. **Direct**: A direct evidence is usually exhibited by TF knockdown or impact on drug response through TF binding or expression.
2. **Regulation of Drug Action**: Target TF has regulatory impact on drug action.
3. **Regulatory Target Direct**: Target TF is a regulator of a gene with direct evidence.
4. **Sibling Drug Evidence**: Target TF has direct evidence for a sibling drug.
5. **Complex Partner Direct**: Target TF is a complex partner of a protein with direct evidence.
6. **Sibling of Direct**: Target TF is a sibling of a protein with direct evidence.
7. **Indirect and No Evidence**: Other types of evidence than the first five are classified as indirect evidence. Indirect evidences are not considered as support for the TF-Drug pair.

1 ELF1 – Rapamycin

Evidence Code: Indirect

Evidence: The TF ELF4 is a part of the mTOR pathway and its expression is regulated by mTOR in CDH8+ cells; rapamycin inhibits mTOR. ELF4 is a paralog of ELF1 (1).

2 ELF1 – Epirubicin

Evidence Code: Direct

Evidence: TF Knockdown (2).

3 MAX – CDDP

Evidence Code: Complex Partner Direct

Evidence: MXD1, a member of the Myc-Max-Mxd family, has been shown to induce CDDP (cisplatin) resistance in hypoxic U 2OS and MG 63 cells (3). Mxd1 (MAD) is a well-known dimerization partner of MAX (4,5), and the reported association of MAX with CDDP is thus likely to reflect the previously demonstrated Mxd1-CDDP relationship.

4 SIN3A – CDDP

Evidence Code: Complex Partner Direct

Evidence: SIN3A is part of the SIN3A-HDAC complex that functions as a repressor and is associated with diseases including cancer (6), and HDAC inhibitors are known to potentiate CDDP activity (7-9).

5 SP1 – Carboplatin

Evidence Code: Direct

Evidence: PMID 24122978 (10).

6 ELF1 – Doxorubicin

Evidence Code: Direct

Evidence: TF knockdown (2)

7 NFYB – 6TG

Evidence: [no TF in database for this drug] NFYB, when induced by E2F1, attenuates apoptosis. Nothing specific to 6TG.

8 MAX – 6TG

Evidence: [no TF in database for this drug] MAX is a heterodimeric partner of c-Myc, which is known to affect mutation rate in the HGPRT gene; the latter in turn is a critical component of 6-TG sensitivity (11).

9 SIN3A – MPA

Evidence: [no TF in database for this drug]

10 ELF1 – Carboplatin

Evidence Code: Regulatory Target Direct; Sibling Drug Evidence

Evidence: ELF1 is one of the top 20 molecular features associated with response to the decitabine:carboplatin combination, as per CTRP database (PharmacoDB). Also, ELF1 is an ETS family transcription factor and other TFs of this family, such as ETS-1, have been shown to be involved in resistance to platinum drugs (12). ELF1 is also believed to regulate BRCA2, which in turn is a key determinant of response to platinum therapy (12).

11 NFYB – 6MP

Evidence Code: No evidence

12 MXI1 – CDDP

Evidence Code: Sibling of Direct

Evidence: MXI1, also known as MXD2, is a member of the MXD1 family, and a closely related member of this family – MXD1 – has been shown, via knock-down assays, to affect CDDP resistance in hypoxic conditions via repression of PTEN (3).

13 YY1 – CDDP

Evidence Code: Direct

Evidence: YY1 knock-down in non-small cell lung cancer (NSCLC) cells increased sensitivity to CDDP (13) though no effect was seen in ovarian cancer cell lines (14).

14 SP1 – CDDP

Evidence Code: Direct

Evidence: SP1 inhibition by tolfenamic acid (TA) was found to increase CDDP response in epithelial ovarian cancer (EOC) cell lines (15).

15 ELF1 – CDDP

Evidence Code: Direct

Evidence: As noted above, ELF1 is believed to regulate BRCA2, which in turn is a key determinant of response to platinum therapy (12). Also, ELF1 binding to DNA was found to increase DNA damage upon cisplatin treatment (16), suggesting that the TF may modulate cisplatin activity.

16 SIN3A – 6TG

Evidence Code: No Evidence

17 PML – Epirubicin

Evidence Code: Sibling Drug Evidence

Evidence: (17) See PML-Doxorubicin association below.

18 REST – Rapamycin

Evidence Code: Regulatory Target Direct

Evidence: A previous study showed that REST exerts regulatory control over the ‘mammalian Target Of Rapamycin’ (mTOR) signaling pathway (18), which is the pathway mediating Rapamycin induced apoptosis. mTOR-Rapamycin is a ‘direct’ evidence pair (19).

19 REST – 6MP

Evidence Code: No Evidence

20 NFYB – Epirubicin

Evidence Code: No Evidence

21 SP1 – Epirubicin

Evidence Code: Sibling Drug Evidence

Evidence: SP1 has been found in various studies to be associated with the activity of Doxorubicin (see below), a drug closely related to Epirubicin (of the anthracycline family). The association between SP1 and Doxorubicin was also reported as significant by our analysis, see below.

22 BHLHE40 – TMZ

Evidence Code: Indirect

Evidence: The apoptosis inhibitor BHLHE40, also known as DEC1, has been shown to be predictive of TMZ response in glioma patients (20).

23 PBX3 – Epirubicin

Evidence Code: No Evidence

24 PML – Doxorubicin

Evidence Code: Direct

Evidence: The promyelocytic leukemia (PML) protein was shown to regulate DNA modification in response to Doxorubicin and knockout of PML was found to abolish Doxorubicin-induced DNA modification and to regulated inhibition of cell proliferation by the drug (21).

25 MXI1 – 6MP

Evidence Code: No Evidence

26 SP1 – Oxaliplatin

Evidence Code: Regulation of Drug Action

Evidence: A study of oxaliplatin response in esophageal cancer cell lines showed that survivin may be a key target of the drug, and that downregulation of survivin by the drug is mediated by SP1, thus arguing in favor of a major role for SP1 in oxaliplatin response (22).

27 MAX – 6MP

Evidence Code: Indirect

Evidence: Evidence Code MAX is a heterodimeric partner of c-Myc, which is known to affect mutation rate in the HGPRT gene (11); the latter in turn is a critical component of 6-MP resistance (23).

28 REST – Carboplatin

Evidence Code: No Evidence

29 NFYB – Doxorubicin

Evidence Code: No Evidence

30 MAX – Epirubicin

Evidence Code: Complex Partner Direct

Evidence: We note that various homo- and hetero-dimers of the MAX family proteins including MAX-MAX and MYC-MAX have similar binding preferences and compete for each other's sites, so our report of an association with MAX may equally well represent an association with MYC-MAX. Another study showed that cMyc, an important binding partner of MAX, attenuates the effect of Doxorubicin (24), a drug closely related to Epirubicin. Also, Doxorubicin was shown to disrupt MYC-MAX interaction (25) and thus affect DNA binding by MYC-MAX, suggesting that regulatory effects of MYC-MAX may underlie Doxorubicin response variation.

31 *ATF2 – Fludarabine*

Evidence Code: Indirect

Evidence: The drug Fludarabine, a purine anti-metabolite, shares properties similar to cladribine and gemcitabine. A gemcitabine combination therapy downregulated ATF2 phosphorylation and activated SAPK/JUNK activation, which in certain cell types yields ATF2-cJun heterodimerization (26). In another study, fludarabine treatment, in a cocktail of therapies, led to the Inhibition of the AKT pro-survival pathway and activation of the JNK-ATF2 pathway; fludarabine therapies seemingly increased the concentration of P-ATF2 (27).

32 BHLHE40 – Epirubicin
Evidence Code: Direct
Evidence: BHLHE40, also known as DEC1, is has been shown to induce cell cycle arrest and premature senescence on Doxorubicin treatment in MCF7 cells (28). The mechanism for this was further explored in (29).

33 SP1 – Doxorubicin
Evidence Code: Regulation of Drug Action
Evidence: SP1 activity is increased in Doxorubicin-resistant HL 60 leukemia cells (30), and SP1 expression levels are predictive of long term prognosis in triple negative breast cancer patients under Doxorubicin treatment (31). Moreover, a recent study (32) showed that insulin pretreatment protects against Doxorubicin induced apoptosis by SP1-mediated transactivation of survivin.

34 MXI1 – Carboplatin
Evidence Code: Indirect
Evidence: MXI1, also known as MXD2, is a member of the MXD1 family, and a closely related member of this family – MXD1 – has been shown, via knock-down assays, to affect cisplatin resistance in hypoxic conditions via repression of PTEN. Cisplatin is a platinum drug closely related to carboplatin, believed to have similar resistance mechanisms (33).

35 ATF2 – Everolimus
Evidence Code: No Evidence

36 PAX5 – Hypoxia
Evidence Code: Indirect
Evidence: Hypoxic conditions increase Pax5 expression in KMS-12BM cells (34).

37 SIN3A – MTX
Evidence Code: Indirect

Evidence: The SIN3A repressor complex is a master regulator of the transcriptional activity of STAT3 (35), and STAT3 has been shown to be directly involved in MTX sensitivity (36).

38 PBX3 – Doxorubicin

Evidence Code: No Evidence

**Supplementary Note 4**: Here we list and discuss the literature evidence we found to show the impact of mediator genes on Drug-induced apoptosis pathway. We looked at the mediators genes identified in the ELF1-Doxorubicin enrichment test.

| Gene | Evidence |
|---|---|
| ANKRD52 | ANKRD52, interacting with PP6, lead to the dephosphorylation of PAK1 (37), which can protect cell from apoptosis through bcl-2 involved pathway (38). |
| ATM | ATM is part of the ATM/TP53 pathway and plays an important role in the activation of TP53 (39). |
| ATP6V0D1 | ATP6V0D1 is a subunit of V-ATPase (NCBI Gene: ATP6V0D1). Inhibition of V-ATPase activity can lead to cell death (40-42). |
| B4GALT2 | In HeLa cells, B4GALT2 is identified as a target gene of p53-mediated apoptosis (43) and its overexpression can induce cell apoptosis (44). |
| BAK1 | |
| COMMD9 | Decreasing the expression of COMMD9 leads to the decreasing of E2F1 activation which enhances p53-mediated apoptosis (45). |
| EIF4A3 | EIF4A3 plays an important role in the alternative splicing of BCL2L1/BCL-X and probably also other apoptotic genes (46). Knocking down of EIF4A3 increases the expression of BCL-Xs in HeLa cells (46). |
| IPO13 | IPO13 mediates the nuclear translocation of PDCD5 (47), which promotes apoptosis through HDAC3/p53 pathway (48). |
| miR6734 | miR6734 up-regulates the expression of p21 and induces cell arrest and apoptosis (49). |
| NABP2 | NABP2 plays an important role in the DNA damage response (50). It can also regulate the stability and transcriptional activity of p53 (51). |
| NEURL4 | NEURL4 regulates the activity of p53 (52). |
| REXO4 | REXO4 is also known as hPMC2, which regulates the expression of p21 (53). |
| TFIP11 | TFIP11 is also known as TFIP-1 (NCBI Gene: TFIP11). Knocking down TFIP induces apoptosis through caspases 3 and caspase 9 (54). |
| TMEM86B | TMEM86B has protein-protein interaction with BCL2L13 (STRING). |
| URI1 | URI1 deficiency is associated with p53-mediated apoptosis (55). |

**SUPPLEMENTARY TABLES**

NOTE: All Supplementary Tables are included as sheets in a separate Excel file. Legends for these tables are provided below.

**Supplemental Table 1**: The number of motifs for each TF. See worksheet “Table 1 Number of Motifs”.

**Supplementary Table 2**: The AUROCs and AUPRCs of TFs in the ASB enrichment using four different TF binding predictors. See worksheet “Table 2 AUROCs and AUPRCs”.

**Supplementary Table 3**: TFs discovered regulating drug response using Delta-MOP alone to predict binding-change SNPs. See worksheet “Table 3 Delta-MOP”.

**Supplementary Table 4**: TFs discovered regulating drug response using Delta-SVM alone to predict binding-change SNPs. See worksheet “Table 4 Delta-gkmSVM”

**Supplementary Table 5**: The results of drug associated enrichment tests for all 888 TF-Drug pairs. See worksheet “Table 5 full results”.

**Supplementary Table 6**: Mediator Genes in ELF1-Doxorubicin Analysis. SNPs in the intersection of TF-Drug hypergeometric tests are: 1) eQTLs associated to Drug Response Genes; 2) ‘binding-change SNPs’ of the TF. DRGs associated with those SNPs are potentially mediators regulated by the TF. Here we list all mediator genes in ELF1-Docorubicin analysis. See worksheet “Table 6 DRGs in ELF1-DOX”.

**Supplementary File 1:** Transcription factor binding motifs.
We obtained 239 PWMs from three different data sources: 1) factor book motifs processed by ENCODE UCSC; 2) motifs downloaded from Factorbook; 3) HOCOMOCO Human (v10) motifs. STAP models are trained separately for each motifs. We excluded the motifs if the maximum prediction of the corresponding STAP model is too small (<0.5). The remaining 225 PWMs, which were used in following analysis, are included in Supplementary File 1.
